# Single-molecule assay for proteolytic susceptibility using centrifuge force microscopy:force-induced destabilization of collagen‘s triple helix

**DOI:** 10.1101/171447

**Authors:** Michael W.H. Kirkness, Nancy R. Forde

## Abstract

Force plays a key role in regulating dynamics of biomolecular structure and interactions, yet techniques are lacking to manipulate and continuously read out this response with high throughput. We present an enzymatic assay for force-dependent accessibility of structure that makes use of a wireless mini-radio centrifuge force microscope (MR.CFM) to provide a real-time readout of kinetics. The microscope is designed for ease of use, fits in a standard centrifuge bucket, and offers high-throughput, video-rate readout of individual proteolytic cleavage events. Proteolysis measurements on thousands of tethered collagen molecules show a load-enhanced trypsin sensitivity, indicating destabilization of the triple helix.

## Introduction

The conventional view of fixed protein structure is evolving towards one of dynamic conformational sampling. This change in perspective requires techniques to interrogate structures that are not static. While the structural technique of NMR spectroscopy offers atomic-level resolution of time-averaged dynamics of small proteins in solution, it lacks the ability to resolve larger structures and complexes. For such systems, transient accessibility of otherwise buried regions can be identified and quantified using enzymatic cleavage assays.^1-3^ These provide significant insight into structural stability, but because analysis requires electrophoretic analysis of products extracted at discrete timepoints, the assays lack real-time monitoring of reaction progress. Furthermore, none of these assays assess the modification of structures by applied force. Force is an increasingly apparent regulator of cellular activity, wherein mechanical actuation of cryptic sites in proteins can control downstream signalling.^4^ There is a need for new methods capable of reading out, in real time, changes in protein conformation resulting from external stimuli such as applied force.

Here, we introduce the technique of high-throughput force-dependent proteolysis using a newly developed mini-radio centrifuge force microscope (MR.CFM), which enables the real-time assessment of molecular structural stability. As proof of concept, we assess the force-dependent modulation of collagen’s triple helical structure.

Collagen is the key component of the extracellular matrix and is the predominant protein in vertebrates, where it confers tensile strength to connective tissues.^5, 6^ Collagen is distinguished at the molecular level by its unique triple helical structure. Not surprisingly given its mechanical importance, there has been substantial interest in understanding how the triple helix deforms when stretched. However, previous studies have reached contradictory conclusions about how the triple helix is modified by applied force (Table 1): does it entropically extend, shear open, unwind, or tighten? These investigations each have limitations: model peptide sequences lack the context of the full-length protein;^6, 7^ triple helical stability can depend on the force field used in simulations;^6, 8, 9^ changes in length of the full protein observed under force do not reveal structural or sequence-dependent information;^10, 11^ and collagenases used to investigate force-induced deformations^7, 12, 13^ are not inert probes because their specialized proteolysis of intact collagen derives from their ability to manipulate its triple helical structure.^14, 15^

**Table 1:** **Reported force-dependent responses of collagen’s triple helix, for forces below 10 pN**

| **<u>Force response</u>** | **<u>Technique used</u>** |
| --- | --- |
| **Stabilizes / tightens** | • force-dependent cleavage with bacterial collagenase<sup>12</sup><br>• steered molecular dynamics<sup>6</sup> |
| **No change (entropically extends)** | • optical tweezers stretching<sup>10</sup><br>• steered molecular dynamics<sup>6</sup><br>• force-dependent cleavage with bacterial collagenase<sup>13</sup> |
| **Destabilizes / unwinds / lengthens** | • force-dependent cleavage with MMP1<sup>7, 13</sup><br>• optical tweezers stretching<sup>11</sup> |
| **Shears open** | • steered molecular dynamics<sup>8</sup> |

To resolve the question of how collagen’s structure is altered by force, we implement the hallmark assay for triple helical stability, trypsin susceptibility,^1^ enhancing its utility by incorporating force into single-molecule proteolysis assays. A tight triple helix blocks proteolytic cleavage by presenting a steric barrier that prevents access of a single polypeptide chain to trypsin’s active site. Thus, only if collagen’s structure locally denatures can cleavage by trypsin occur.^1^ If a stretching force destabilizes the triple helix, an increase in the proteolysis rate should be observed (Figure 1). Conversely, if no change in structure occurs or if the helix tightens, the cleavage rate should remain unchanged or be reduced. To assess the effect of force on collagen’s stability, we use MR.CFM to probe thousands of individual collagen molecules and determine the how force affects their rate of proteolysis by trypsin.

**Figure 1.**
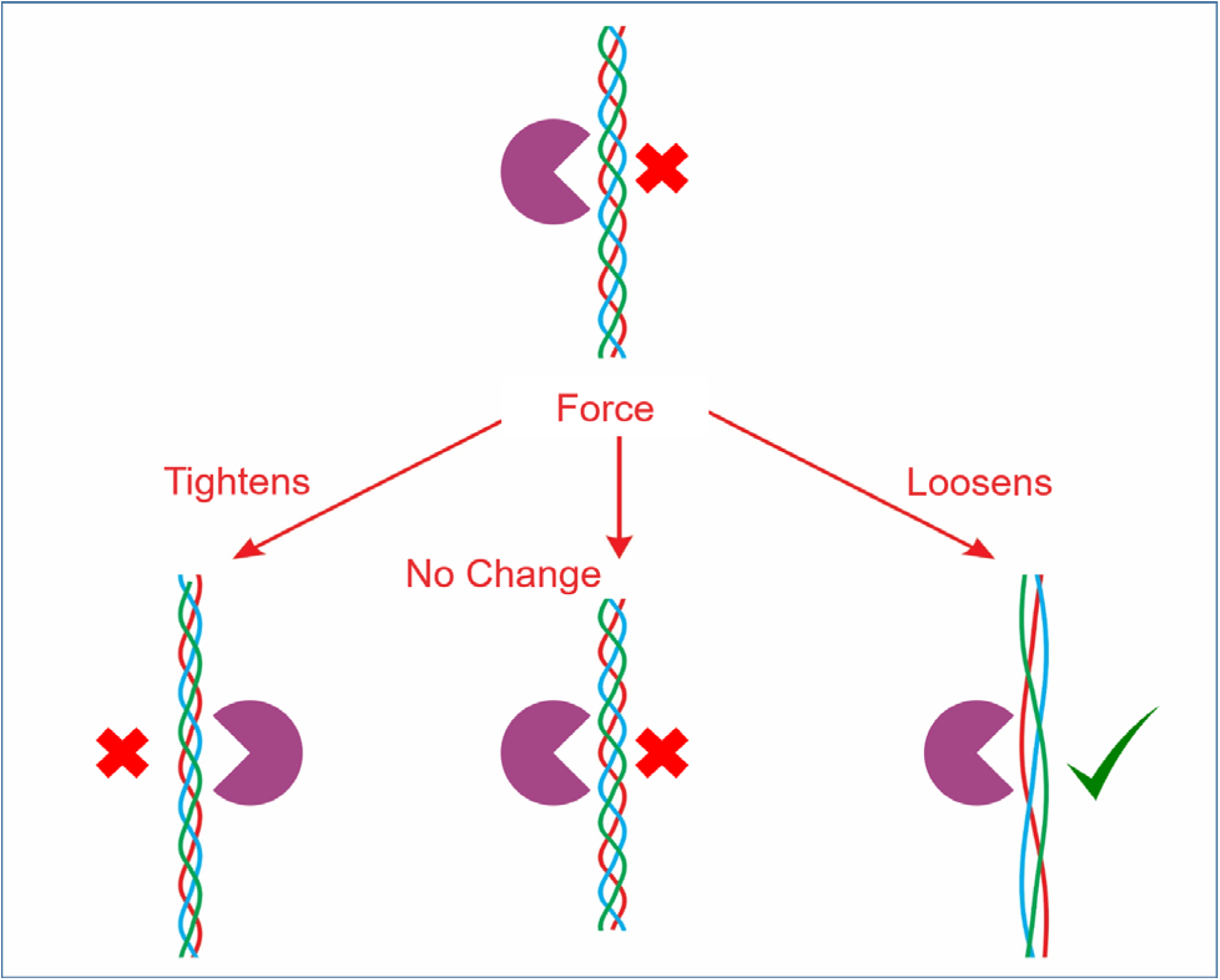
Possible responses of collagen’s triple helix to applied stress. In the absence of applied force, a stable triple helix is resistant to proteolysis by trypsin (top). If the helix tightens or remains unchanged by force, it remains resistant to cleavage by trypsin. Only if force induces a destabilization of the helix will proteolysis by trypsin be possible (lower right).

## Results and Discussion

### MR.CFM – Mini-Radio Centrifuge Force Microscope

To perform high-throughput proteolysis assays in a force-dependent manner, we developed a new instrument dubbed the mini-radio centrifuge force microscope (MR.CFM, Figure 2). Centrifuge force microscopy requires only a microscope capable of stable imaging at high acceleration and dense particles on which to exert force, and can have a force range orders of magnitude larger than magnetic or optical tweezers.^16^ Our instrument was designed to be easily adopted by biological research labs, as it is a “plug-and-play”, modular, entirely wireless brightfield microscope that can be placed into the bucket of a conventional benchtop centrifuge, counterbalanced simply by tubes of water. The force is determined by the rotational frequency of the centrifuge, ω (typically hundreds-thousands of RPM), by the distance between the sample and the axis of rotation *r* (typically tens of centimetres) and by the mass of bead (*m*, relative to the displaced water) tethered to the molecule of interest: *F*=*m*ω^2^*r*. Unlike the limited region of constant force in alternative single-molecule force techniques,^17^ the CFM provides a controllable, uniform force across the *entire* rotating sample chamber.

**Figure 2.**
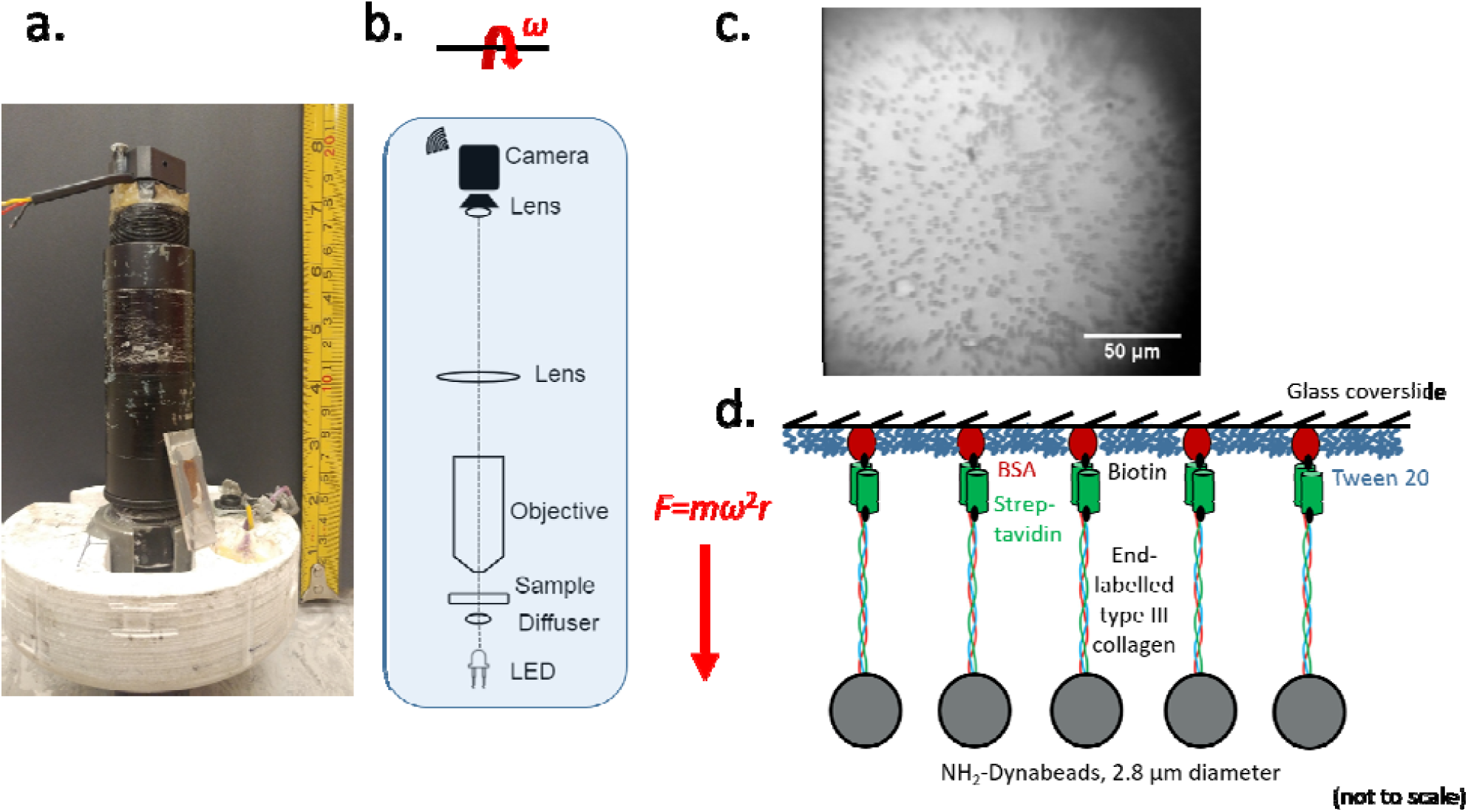
Mini-Radio Centrifuge Force Microscope (MR.CFM) and collagen tethering. a) Photograph of the microscope, with the centrifuge bucket insert at the bottom and a sample chamber leaning at the side. The height of the assembly is less than 16 cm and its mass is 421 g. b) Schematic of the optical elements within the microscope. A force *F*=*m* ω^2^*r* is exerted on all beads of relative mass *m* within the sample chamber, located a distance *r* from the central rotation axis, when the rotor spins at angular frequency ω. c) Image of microspheres subjected to 9 pN of force in MR.CFM, tethered to the surface by collagen molecules. Although each bead occupies only a few pixels in the final image, they are easily detected. d) Schematic of tethering geometry. Single molecules of collagen are tethered to a glass slide via biotin-streptavidin linkages, and are covalently linked to the surface of heavy beads via thiol-amine coupling.

The layout of MR.CFM was driven by optical simplicity and low mass. Thus, in contrast to alternative approaches,^18, 19^ our instrument has a straight optical path and requires no mirrors. Besides ease of optical design, the straight layout positions the camera closer to the centre of rotation, subjecting it to less force during operation. In addition to substantially reduced cost, MR.CFM affords significant advantages over previous compact designs, by (a) utilizing an unmodified centrifuge, thereby being amenable to use in shared facilities;^19^ and (b) offering continuous radio-frequency video output, permitting unlimited video-rate capture and digitization.^18^ The latter advantage confers the ability to read out molecular kinetics over experimental timescales of hours, without the need to sacrifice temporal resolution.

With a CFM, throughput is limited only by the ability to resolve individual tethered particles. In our instrument, image quality is limited by the camera, which was selected to provide exceptional value for financial and weight budgets: the image quality is sufficient for bead detection and the “binary” decision about the presence or absence of a bead (see below); the analog radio-frequency signal can be captured and digitized at video rates in real time; the cost of the camera is only $25 USD at the time of this writing; and its mass is only 9 grams. Moreover, because each camera unit has a unique RF output, further multiplexing of experiments would involve simply cloning MR.CFM with a different camera unit, and capturing video streams to the computer from parallel experiments within one centrifuge.

A wide field of view and high-efficiency surface chemistry enable the simultaneous visualization of hundreds of particles. High-throughput analysis is enabled by manipulating the raw image stream acquired by MR.CFM and computationally detecting and counting scores of particles (Figure 3). The image processing used in this work minimizes user input and is implemented within a semi-automated ImageJ macro. With the use of this macro, beads occupying only a few pixels in the wide-field images from MR.CFM can be easily detected.

**Figure 3.**
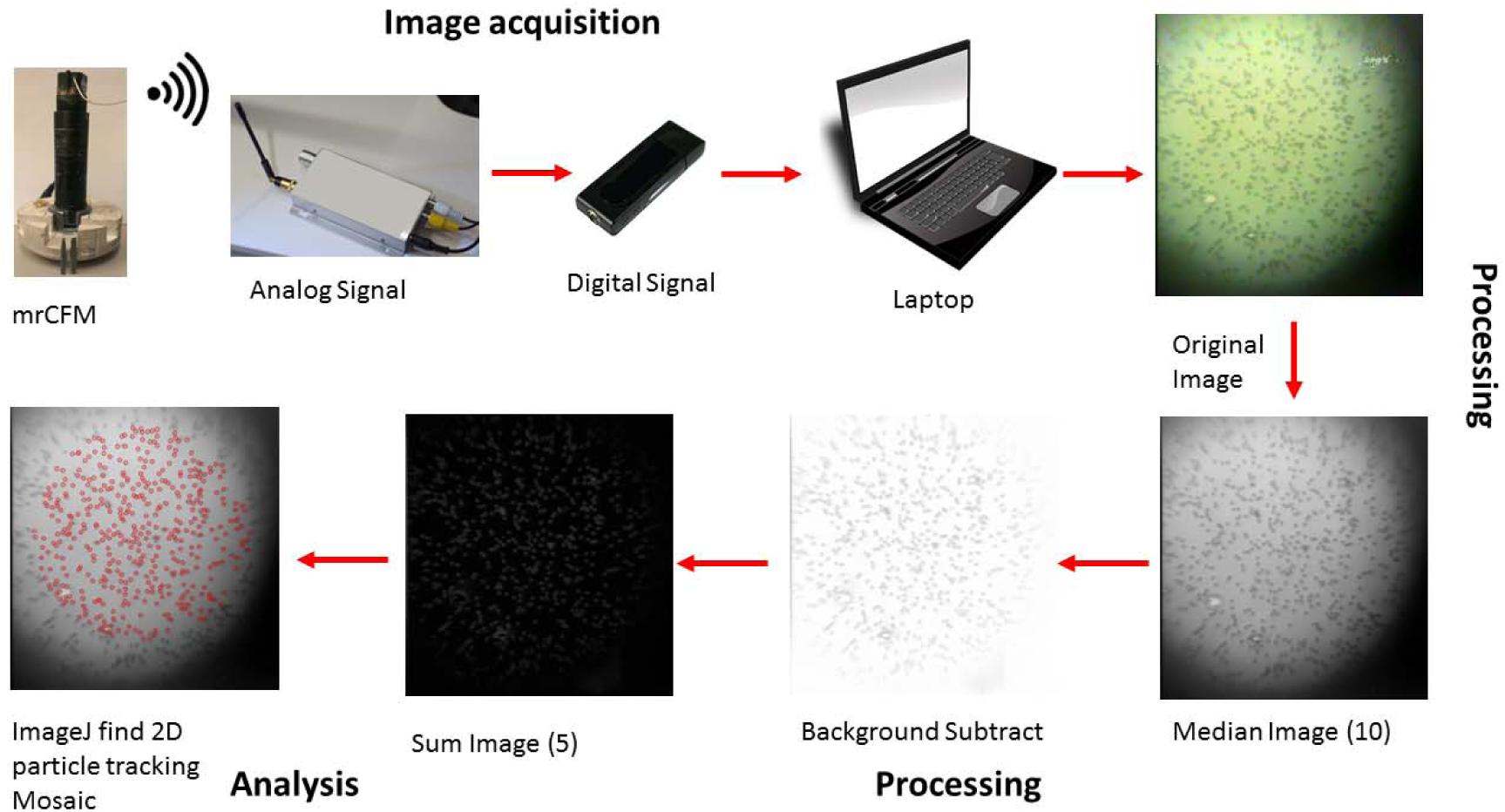
Work flow for MR.CFM acquisition, processing and analysis of bead images. Images are acquired with the wireless camera in MR.CFM, then transmitted via radio signal to the AV receiver. The AV receiver collects the analog signal, which is output to a USB analog-to-digital converter attached to a laptop computer. Arcsoft Totalmedia 3.5 software is used to provide real-time visualization and simultaneous storage of the image stream. The original output image files from the MR.CFM (shown here at 9 pN of force) are processed in ImageJ to improve contrast and remove rotation-speed dependent interference. Particles are counted using the Mosaic particle tracking program. The resultant overlaid image visually shows that this procedure is robust and identifies beads within the defined search area.

### Collagen proteolysis under force

To determine whether tension affects stability of collagen’s triple helix, we determined how an applied force alters its rate of cleavage by trypsin. Single type III collagen molecules were tethered by a heavy microsphere to a surface, and force was applied from gravity (*F* = 0.067 pN ≈ 0) or by force from rotation of a centrifuge (Figure 2d). Tether rupture kinetics were measured in the presence and absence of trypsin. Supplementary Video 1 provides an example experimental data set with collagen strained by 9 pN of force, in the presence of trypsin.

From measurements on thousands of distinct single-molecule tethers, our data show a significantly enhanced rate of collagen cleavage by trypsin in the presence of applied force (Figure 4 and Supplementary Figure 1). Decay curves are well described by single-exponential functions, which were fit to the data. We found the trypsin-dependent cleavage rate to increase from k_Tr,0_ = 0.007 ± 0.013 min^-1^ to *k*_Tr,F_ = 0.227 ± 0.037 min^-1^ as the force increased from 0 to 9 pN. The enhancement of proteolysis by a relatively low level of force implies that collagen’s triple helix destabilizes when stretched, increasing the accessibility of otherwise sterically hindered chains to the protease active site.

**Figure 4.**
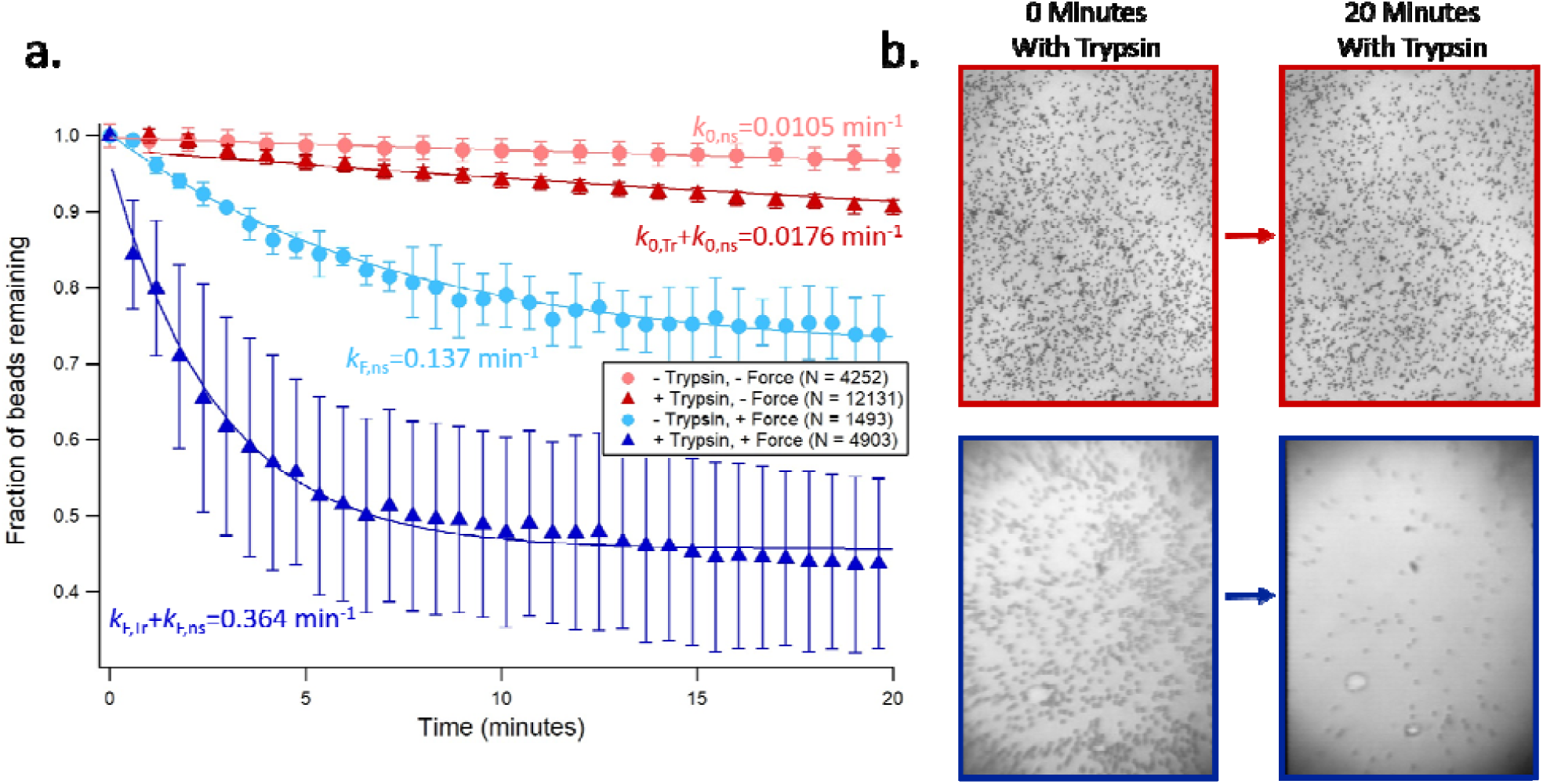
Collagen proteolysis by trypsin is enhanced by force. a) Collagen tethers rupture at higher rates in the presence of force and trypsin (dark blue triangles) than in the absence of either, implying their force-induced denaturation. Subsampled data of the fraction of beads remaining are shown for clarity, with full data sets available in the supporting information. Fits and error bars are described in Supplementary Note 3. b) Representative images of collagen-tethered beads at the beginning (left) and after 20 minutes (right) of incubation with trypsin. Top no force (67 fN; gravitational force on beads), recorded using a conventional bright-field microscope. Bottom: 8.8 pN of force, recorded within MR.CFM.

The loss of tethered beads from the surface, even in the absence of enzyme, does not occur with different surface chemistry (Supplementary Figure 2) so may be attributed to mechanically weak interactions, perhaps between biotinylated BSA and silane,^20^ rather than from shear-induced rupture of the triple helix.^8^ Enzyme-independent detachments were accounted for in fitting the trypsin cleavage data. In addition, the observation of a non-zero cleavage rate at zero force is consistent with the expectation that type III collagen is weakly susceptible to trypsin digestion in solution at room temperature.^21^

Our observations of force-induced weakening of the triple helix are consistent with force-enhanced collagen cleavage observed using the collagenase matrix-metalloprotease-1 (MMP1).^7,13^ Interestingly, of the 81 possible trypsin sites per type III collagen chain, previous analysis of collagen fragments produced by trypsin proteolysis identified one primary cleavage site,^21^ located in a “loose” region of the triple helix that is shared by the unique recognition sequence of MMPs.^22^ The observed force-enhanced proteolysis of collagen by trypsin, as found for MMP1, may be a result of both enzymes probing the same region of the collagen triple helix. In contrast to MMP1, trypsin has not evolved to interact with triple helical collagen, and thus may be considered a more passive probe of externally-induced changes in collagen’s structure.

Just as distinct sequences of triple helix can possess different thermal stabilities and helical pitch,^5^ it is possible that collagen has a molecular structure whose response to applied force is modulated by its local sequence. Future experiments could examine this question with multiple proteases targeting distinct regions of collagen’s sequence, thereby mapping the sequence-dependent mechanical stability within the native, full-length protein. How distinct mechanical signatures along the length of each collagen interact with and modulate biochemical and cell biological interactions is an exciting question to pose for future studies. More generally, the real-time, highly multiplexed approach of MR.CFM enables the discovery and characterization of transiently accessible and mechanically cryptic sites within native states of proteins,^23^ protein-DNA structures and protein assemblies.

## Methods

### Mini-Radio Centrifuge Force Microscope (MR.CFM)

MR.CFM is a wireless, fully self-contained, compact CFM that exploits the rotation control of a commercial benchtop centrifuge. MR.CFM fits within a single bucket of a Beckman Coulter Allegra X-12R centrifuge, equipped with an SX4750A ARIES “Swinging Bucket” Rotor Assembly. This assembly has an imbalance tolerance of ±50 g. MR.CFM was built using as many commercial parts as possible for ease of reproduction. The final height and weight for MR.CFM are 154 mm and 421 g (including batteries), respectively. These fall well within the length and weight budgets of the centrifuge rotor.

Schematics and parts lists are provided in Supplementary Figure 3 and Supplementary Table 1, with a description of the design provided in Supplementary Note 1. Drawings of the few custom parts are provided in Supplementary Figure 4 and Supplementary CAD drawings. Note that, while the designs provided are specific to this rotor assembly, MR.CFM can easily be adapted for other benchtop centrifuges, primarily by modifying the base.

### Image acquisition and particle detection

The image analysis workflow to acquire, process and analyze data from the raw image stream is shown schematically in Figure 3 and is described in Supplementary Note 2. The ImageJ macro used in image processing is provided as Supplementary material. Particles are counted with the Mosaic particle tracker 2D/3D plugin for ImageJ.^24^

### MR.CFM sample chambers

Glass slides and coverslips were cut to 12 mm by 33 mm to fit the sample holder. Sample chambers were created between a cover slip (borosilicate glass No. 1; VWR, 48393 106) and glass slide (soda lime glass 2 mm, Logitech Limited) by strips of glue (JB Weld Original), with a depth of approximately 300 µm.

### Surface chemistry

We implemented a blocking and specific tethering system that relies on a self-assembled monolayer (SAM) of Tween-20 for blocking and on biotinylated BSA for specific tethering. Modifications to the published protocol for surface blocking^20^ are provided in Supplementary Note 3.

### Collagen functionalization

Type III collagen was used in these experiments due to its chemically accessible C-terminal cysteine residues, unique among fibrillar collagens. Recombinant human type III collagen (Fibrogen FG-5016) was end-labelled for manipulation as previously described,^25^ with information provided in Supplementary Note 3. Briefly, this approach utilized biotinylation of introduced N-terminal aldehydes the covalent linking of C-terminal thiols to amine-functionalized superparamagnetic microspheres (2.8 µm diameter, M270 Dynabeads).

### Single-Molecule Proteolysis Experiments

All trypsin cleavage experiments were performed at room temperature with the following final concentrations: trypsin (Sigma T1426) 2.0 mg/ml, 1X reaction buffer (0.1 M Tris, 0.4 M NaCl, pH 7.4), and collagenated beads (< 10^8^ beads/ml). To minimize room-temperature proteolysis of collagen by trypsin (Supplementary Figure 5), all solutions were kept at 4°C before use. Experiments under 9 pN of force were performed in MR.CFM, while zero-force (67 fN) experiments utilized a bright-field microscope (Olympus BX51, equipped with 10X Plan N objective). Detailed experimental protocols are provided in Supplementary Note 3, as is information about the number of distinct experimental runs for each condition. In the absence of collagen, labelled beads remained tethered to the surface in the presence of trypsin (Supplementary Figure 6), demonstrating that the trypsin-dependent removal of collagen-labelled beads resulted from collagenolysis.

The number of potential trypsin cleavage sites in type III collagen was given by the Expasy PeptideCutter tool,^26^ querying from sites 154-1221 of UniProtKB entry P02461. This sequence represents the collagen domain of human type III procollagen.

### Data Analysis

The fractions of beads remaining as a function of time for both trypsin- and trypsin-independent experiments were well described by single-exponential decay processes:

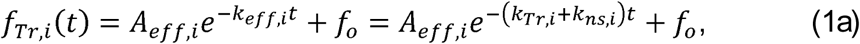

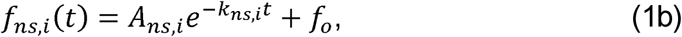

where the index i=0 for the no-force experiments and i=F for experiments under 9 pN of force. *f*_o_ represents the fraction of beads that do not detach in each experimental data set. “Tr” refers to trypsin-dependent rates, while “ns” refers to “non-specific” bead detachment in the absence of enzyme. Because both trypsin-dependent cleavage and nonspecific rupture of tethers contribute to the observed events, trypsin-dependent data were described by an effective rate constant, *k*_eff_, which contains contributions from both processes. This arises directly from the following first-order decay model:

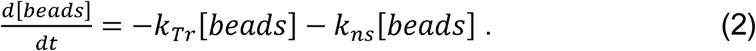

Fitting of the decay data with equations (1) was performed in IGOR Pro (WaveMetrics, Inc.), weighted by the errors associated with each point (Supplementary Note 3). Fit parameters are presented as best fit values +/- standard deviation. Rate constants obtained from fitting each data set are reported in Supplementary Table 2. To determine whether force enhances or hinders collagen proteolysis, we compared the trypsin-dependent rates *k*_Tr,F_ and *k*_Tr,0_.

### Data availability

The datasets generated and analysed during the current study are available from the corresponding author on request.

## Acknowledgements

This research was funded by a Discovery Grant from the Natural Sciences and Engineering Research Council of Canada (NSERC). We gratefully acknowledge Aaron Lyons for AFM imaging and quantification, Wesley Wong and Ken Halvorsen for helpful discussions, and David Lee for the use of the bright-field microscope. We thank John Bechhoefer, Edgar Young, and members of the Forde laboratory for critical readings of the manuscript.

## Author contributions

MWHK and NRF designed the research. MWHK built MR.CFM and performed all experiments. MWHK and NRF analysed the results and wrote the manuscript.

